# Baseline epidemiology and associated dog ecology study towards stepwise elimination of rabies in Kwara state, Nigeria

**DOI:** 10.1101/2020.06.08.140517

**Authors:** Ahmad Ibrahim Al-Mustapha, Abubakar Ahmed Tijani, Oyewo Muftau, Folashade Onatola Bamidele, Ahmed Ibrahim, Muhammad Shuaib Osu, Babasola Olugasa, Muhammad Shakir Balogun, Grace Kia, Stella Mazeri, Annamari Heikinheimo

## Abstract

Understanding domestic dog population dynamics and ecology is necessary for any effective rabies control program. This study was conducted as part of the baseline epidemiological studies necessary for the establishment of the Kwara Rabies Rapid Alert System “KRRAS”. The aim of this study was to determine the dog population structure of Kwara State by assessing the dog ownership, vaccination status, and prevalence of dog bites.

A total of 1,460 questionnaires were administered to respondents in the three senatorial zones of the state using Open Data Kit (ODK) between June 2019 to January 2020.

Of the 1460 households surveyed, 293 (20.1%) owned at least one dog with an average of 2.25 dogs per household. The male to female ratio was 2.2:1 and 85% (n=250/293) of the owned dogs were local breeds. A total of 785 dogs was enumerated (659 dogs from 293 households and 126 free-roaming dogs) and 7811 persons which resulted in a dog-human ratio of 1:9.95. The estimated dog population is 376,789 (95% CI: 343,700 – 379,878). The dog anti-rabies vaccination coverage was 31% (n=92/293). The prevalence of dog-bite was 13% (n=193/1460) of which only 27% of the victims (n=61/225) received post-exposure prophylaxis (PEP). The ethnicity of respondents had a significant impact on dog ownership. Yoruba’s more often (OR: 2.2-5; 95% CI: 1.2 − 12.4; p < 0.001) owned dogs than other tribes. The vaccination status of owned dogs was greatly impacted by the level of education of the respondents (OR: 5.03; 95% CI: 1.5073 − 16.8324; p *<* 0.001); the breed of the dog with exotic dogs being more vaccinated (OR: 2.8; 95% CI: 0.7150 − 10.857; p *<* 0.001) and the confinement of the dog (OR: 2.07; 95% CI: 1.1592 – 3.7037; p < 0.001). Always confined dogs were twice more vaccinated than non-confined dogs. The results of the study showed that the vaccination coverage needs to be increased, the number of dog bites needs to be reduced, the number of non-confined dogs needs to be reduced and stray dog control strategies need to be implemented. The findings of this study showed very low vaccination coverage for dogs which is below the 70-80% target recommended for herd immunity by the world health organization.

## Background

Rabies is one of the oldest and most terrifying diseases known to man (Horton et al., 2015). It is a fatal viral disease caused by one of the seven lineages of *lyssavirus* with a distinct “bullet” shape belonging to the family *Rhabdoviridae* (Badrane et al., 2001). Domestic dogs account for over 99% of human death from rabies (WHO, 2013).

It causes approximately 59,000 (95% CI: 25 − 159,000) human deaths annually mainly in Asia and Africa with 40% of people bitten by suspect rabid animals are children under 15 years of age (OIE, 2017a). Annually, it is responsible for 3.7 million (95% CI: 1.6 − 10.4 million) disability-adjusted life years (DALYs) and 8.6 billion USD (95% CI: 2.9 − 23.5 billion) economic losses (Hampson et al., 2015).

Rabies is endemic in Nigeria and has remained one of the most important neglected diseases of public health concern in the country. Despite the under-reporting of dog-bites incidents, about 10,000 cases of dog-bites are reported annually (NCDC, 2018). The rabies viral antigen has been detected in the brain tissues of apparently healthy dogs slaughtered for human consumption (Suleiman et al., 2020; Mohammed et al., 2019; Kia et al., 2018; Hambolu et al., 2014; Garba et al., 2010).

Several factors such as poor vaccination strategy, lack of sufficient vaccines, presence of stray (community) dogs and illegal trade of dogs within and across countries are major contributory factors to the endemicity and transboundary movement of rabies in Nigeria (Kia et al., 2018; Ogo et al., 2011; Ogunkoya et al., 2008). The epidemiology of rabies has been described in Nigeria with dogs as the main reservoir of the disease (Kia et al., 2018; Garba et al., 2011; Ameh et al., 2014). Most industrialized countries have eliminated rabies from domestic dog populations. However, in the majority of developing countries, rabies remains endemic in domestic dog populations and poorly controlled (Coleman et al., 2004). The presence of unvaccinated free-roaming dogs (FRD) amidst human settlements is a major contributor to the high incidence and maintenance of rabies in Nigeria.

Mass dog vaccination is the most cost-effective strategy for preventing human rabies (OIE, 2017b). Vaccinations can be by parenteral administration of inactivated vaccines or by the use of baits for oral vaccination of dogs (Maki et al., 2017). Dog vaccination reduces deaths attributable to rabies and the need for post-exposure prophylaxis (PEP) as a part of dog bite patient care (Cliquet et al., 2018).

Through the implementation of the *Global strategic plan to end human deaths from dog-mediated rabies by 2030* (WHO, 2017), the importance of projects such as KRRAS cannot be over-emphasized. The project is divided into three phases: Baseline epidemiological assessment studies; Mass dog vaccinations; Surveillance of dog-bite cases in the state using the existing disease surveillance and notification officers (DSNOs) in the state. This study, as well as three others, are the epidemiological basis for the proposed Kwara Rabies Rapid Alert System (KRRAS)-a one health system designed to improve reporting of dog-bite cases and enhance diagnosis of rabies.

Our baseline data aims to generate evidence-based data that can be used for the dog leash law (in Kwara State). Estimation of the dog population and ecology in the state is important in vaccine procurement and prioritizing intervention plans.

## Methods

### Study area

Kwara State, located in the southern guinea savannah zone of Nigeria between latitude 8.9669°N, and longitude 4.3874° E. The state has a population of 3,599,800 (NPC, 2020). It is located in Northcentral Nigeria. The state has three agro-ecological and geographical zones (Northern, Central, and Southern) with vast land and with varying climatic conditions.

### Period and course of the survey

A cross-sectional survey was carried out to generate a baseline dog population, dog to human ratio, anti-rabies and DHLPP vaccination coverage as well as assess the incidence of dog bites in Kwara state (n=1460). We developed a structured questionnaire that was used for transect walk (House to house and street counts of dogs) in selected areas. The questionnaire was pre-tested in a pilot study and adjusted accordingly before being used in this survey. The questionnaire is composed of three sections: a) Owner demographics b) Dog ownership and vaccination status c) Incidence of dog bites and its management.

The survey was conducted from June 2019 to January 2020 in the three senatorial zones of Kwara state. A multi-stage sampling (Kwara state → 3 senatorial zones → local government areas → communities → areas → streets → households) of the respondents was carried out from all the communities in the state. Furthermore, a systematic random sampling of households was conducted to select respondents in each street. A sampling interval of 5 houses was used for this study.

### Organization of survey team and movement plan

The Kwara Rabies Research Team (Kw-RRT), supported by a team of volunteers, composed of field epidemiologists, Unilorin One Health Students, and data collectors carried out the survey. They were trained on the methodology of the survey and were divided into five groups of two members, plus a group leader. They were randomly assigned separate predetermined routes. The survey was conducted on each chosen street. Starting from the first houses on the right side, every 6^th^ house was selected and an adult member (18 years and above) was interviewed using the open data kit application (available on mobile platforms). In areas where this was not applicable due to the lack of organized road connections as commonly seen in semi-urban and rural areas, the polio-vaccination micro-plan was adopted, using the polio house markings as a guide.

### Free Roaming Dog Enumeration Technique

We used Beck's definition of a free-roaming dog (FRD) as “Any dog observed without human supervision on public property or private property with immediate unrestrained access to public property” (Berman and Dunbar, 1983).

Using the photographic recapture technique (Beck, 1983); we used photography to prevent counting a dog twice within the same area. Surveys were alternated between mornings and late evenings on two alternate days. On each day of the study, counting of dogs was carried out in the morning (between 6 and 9 a.m.) and in the evening (5.30 to 7.00 p.m.). Every dog within a 5m radius was photographed and recorded.

The number of counted dogs in the selected streets in each area was estimated using the formula:

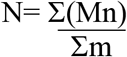

Where N= the estimate of the dog population of Kwara state.

M= the number of observed/photographed dogs each time.

m= the number of dogs identified as previously observed.

n= Each days’ observation (M) – previously observed dogs (m).

### Data analysis

Data were retrieved from the African Field Epidemiology Network (AFENET) server and analysis were performed by Chi-square and logistic regression using Minitab 19.1.1 (Pennsylvania, USA).

### Development of logistic regression models

A bivariate logistic regression analysis was used to evaluate the effects of factors affecting dog ownership and vaccination status. Furthermore, a multivariate logistic regression model (using logit function) was fitted to analyze statistically significant factors affecting dog ownership and their vaccination status.

## RESULTS

### Demography of Respondents

The survey was conducted in the three senatorial zones (North, Central, and South) of the state. The total number of respondents was 1,460. Of these, 72% (n=1045/1460) had either secondary or tertiary education which made administering the survey in English feasible. However, the questionnaire was administered in the local language of the community when needed. Tradesmen/Artisans accounted for over 47% of the respondents (Table 1).

**Table 1.** Demographic structure of respondents in the three senatorial zones of Kwara state.

| Senatorial Zone | No of respondents (%) |
| --- | --- |
| Kwara South | 364 (25) |
| Kwara North | 562 (38.5) |
| Kwara Central | 534 (36.5) |
| Total | 1460 (100%) |
| Sex |  |
| Female | 382 (26.1) |
| Male | 1072 (73.5) |
| Prefer not to say | 6 (0.4) |
| Total | 1460 (100%) |
| Religion |  |
| Christianity | 523 (35.8) |
| Islam | 914 (62.6) |
| others | 3 |
| Prefer not to say | 21(1.4) |
| Total | 1460 (100) |
| Level of education |  |
| No western education | 187 (13.5) |
| Primary | 165 (11.9) |
| Secondary | 581 (41.9) |
| Tertiary | 424 (30.6) |
| Prefer not to say | 73 (5) |
| Other | 30 (2.1) |
| Total | 1460 |
| Occupation |  |
| Artisan | 286 (20.4) |
| Civil servant | 128 (9.1) |
| Driver | 20 (1.4) |
| Farmer | 169 (12) |
| Fisherman | 3 (0.2) |
| Laborer | 12 (0.9) |
| Other professions | 64 (4.6) |
| Security officer | 17 (1.2) |
| Student | 253 (18) |
| Trader/ businessman | 376 (26.9) |
| Unemployed | 72 (5.2) |
| Prefer not to say | 60 (4.1) |
| Total | 1460 |

### Dog Ecology

A total of 126 free-roaming dogs (FRDs) were counted during the survey across the three senatorial zones. However, a total of 659 dogs were owned by 293 respondents resulting in an average of 2.25 dogs per visited household. During the survey, 7811 people were counted and a total of 785 dogs (FRD and owned) were counted giving a dog to human ratio of 1:9.95. Therefore, with a total population of 3,599,800; we estimate a total dog population of 361,789 (95 CI: 343,700 – 379,878) dogs in Kwara state (Table 2). The descriptive statistics of owned dogs are provided in Table 3. The local breeds of dogs (Mongrel) accounted for 85 of owned dogs. Most respondents (52%, n=198/380) kept dogs for security whereas hunting dogs accounted for another 28% (n=105/380) of owned dogs. Based on the use of the dog, 69% (n=202/302) partially/ never confine their dogs. Dogs aged between 1-3 years were the most reported (56%) with 92% of all owned dogs being 5 years or younger.

**Table 2.** Dog and human population structure in baseline rabies survey of Kwara state.

| Variables |  |
| --- | --- |
| Number of households sampled | 1460 |
| Total number of people | 7811 |
| Total number of dogs (Owned + FRDs) | 785 |
| Dog human ratio | 1:9.95 |
| The estimate of dog population in Kwara State | 361,789 |

**Table 3.** Characteristics of owned dog populations in Kwara state.

| Dogs in your household? | Number of respondents (%) |
| --- | --- |
| Yes | 293 (20.1) |
| No | 1167 (79.9) |
| Total | 1460 (100) |
| Gender of dog |  |
| Female | 92 (31) |
| Male | 201 (69) |
| Total | 293 (100) |
| Breed of dog |  |
| Cross-bred | 27 (9.3) |
| Exotic | 16 (5.4) |
| Local | 250 (85.3) |
| Total | 293 (100) |
| Age of dogs |  |
| < 1 year | 53 (18) |
| 1-3 years | 164 (56) |
| 3-5 years | 54 (18) |
| > 5 years | 22 (8) |
| Total | 293 (100) |
| Use of dog |  |
| Breeding | 31 (8) |
| Herding | 5 (1) |
| Hunting | 105 (28) |
| Pet | 41 (11) |
| Security | 198 (52) |
| Total | *380(100) |
| Confinement of dog |  |
| Always | 80 (27) |
| Never | 152 (52) |
| Partial | 50 (17) |
| I don't know | 11 (4) |
| Total | 293 (100) |

\*Some dogs were used for more than one purpose.

### Dog vaccination status

Of the 293 households surveyed, only 31% (n=92/293) were previously vaccinated against rabies. Among the dogs vaccinated against rabies, only 22% (n=20/92) had received DHLPP^®^(a recommended pentavalent vaccine for puppies at 8 weeks of age meant to provide immunity against Canine distemper, Hepatitis, Leptospirosis, Parainfluenza, and Parvovirus). With 52% (n=48/92) of all vaccinations administered at home (Table 4).

**Table 4.** Evaluation of the vaccination status of owned dogs in Kwara State.

| Vaccination against rabies | Number of respondent households (%) |
| --- | --- |
| Yes | 92 (31) |
| No | 201 (69) |
| Total | **293 (100)** |
| Vaccination against DHLPP® |  |
| Yes | 20 (22) |
| No | 72 (78) |
| Total | **92 (100)** |
| Where vaccination against rabies was received |  |
| At home | 48 (52) |
| Private Veterinary Clinic | 27 (30) |
| Gov't Vet clinic | 17 (18) |
| Total | **92 (100)** |
| Pet green book |  |
| Yes | 24 (26) |
| No | 68 (74) |
| Total | **92 (100)** |
| Age of first vaccination |  |
| 3-6 months | 14 (15) |
| 6-12 months | 12 (13) |
| 1-2 years | 24 (26) |
| 2-3 years | 13 (14) |
| 3-4 years | 6 (7) |
| 4-5 years | 12 (13) |
| 5 years + | 11 (12) |
| Total | **92 (100)** |
| Modes of dog depopulation |  |
| Give away | 36 (55) |
| I Keep my pups | 10 (15) |
| Killed | 4 (6) |
| Sold | 15 (23) |
| Total | **65 (100)** |

### Incidence and management of dog bites in Kwara state

Of the 1460 respondents, 13% (n=193/1460) had a history of dog-bite with 86% being beaten once (63%, n=154/246) or twice (23%, n=56/246) as presented in Table 5. About 13% (n=27/225) and 32% (n=72/225) use antibiotics and traditional methods (herbs) to treat dog-bite wounds (Table 5).

**Table 5.** Evaluation of dog-bite incidence and it’s management.

| Ever bitten by a dog? | No. of respondents (%) |
| --- | --- |
| No | 1267 (87) |
| Yes | 193 (13) |
| Total | 1460 (100) |
| No of bites incidents |  |
| 1 | 154 (63) |
| 2 | 56 (23) |
| 3 | 21 (9) |
| 5 | 15 (6) |
| Total | 246 (100) |
| Outcome of dog |  |
| Alive | 72 (36) |
| Died | 11 (6) |
| I don't know | 12 (6) |
| Killed | 59 (30) |
| Ran away | 46 (23) |
| Total | 199 (100) |
| Treatment given to the dog-bite victim |  |
| Antibiotics | 27 (13) |
| Non-specific | 64 (28) |
| PEP | 61 (27) |
| Traditional | 73 (32) |
| Total | 225 (100) |

### Bivariate logistic regression analysis of factors affecting dog ownership and vaccination status in Kwara State

In the univariate analysis for dog ownership, there was no evidence that religion had an impact on dog ownership in Kwara state (Table s1). Based on ethnicity, Yoruba people more often (OR: 2.2-5; 95% CI: 1.2-12.4; p < 0.001) owned dogs than other tribes (Table s1). In the multivariate analysis, the odds of having a dog (OR: 1.8-4.4; 95 CI: 0.5-11.3; p < 0.001) was still higher if the respondent was Yoruba after adjusting for the occupation and religion (Table S2).

In the univariate analysis for vaccination status, Male dogs were 1.7x (95% CI: 0.97 – 3; p = 0.063) more likely to be vaccinated than bitches. Previously bitten persons are 1.43x (95% CI: 0.8-2.5; p = 0.2) more likely to have vaccinated their dogs (Table 6).

**Table 6.** Univariate analysis of variables affecting owned dogs in Kwara State.

| Outcome variable | Variable | Referent |  | Odds Ratio (95 CI) |  | X<sup>2</sup> | DF | p-value |
| --- | --- | --- | --- | --- | --- | --- | --- | --- |
| Vaccination status of owned dogs | Gender of dog | Female | Male | 1.7 | (0.971 - 2.977) | 3.45 | 1 | 0.063 |
|  | Confinement of dogs | Always | Never | 0.48 | (0.2700 - 0.8626) | 7.94 | 3 | 0.047 |
|  |  |  | Partial | 1.00 | (0.4861 - 2.0570) |  |  |  |
|  |  |  | I don't know | 0.56 | (0.1387 - 2.2817) |  |  |  |
|  | Breed | Cross-bred | Exotic | 2.79 | (0.7150 - 10.857) | 18.42 | 2 | <0.0001 |
|  |  |  | Local | 0.33 | (0.1488 - 0.7454) |  |  |  |
|  | Previous dog-bite incident | No | Yes | 1.43 | (0.8279 - 2.4628) | 1.64 | 1 | 0.2 |
|  | Use of dog | Breeding | Herding | 4.33 | (0.4227 - 44.4283) | 1.88 | 4 | 0.757 |
|  |  |  | Hunting | 1.96 | (0.5045 - 7.6906) |  |  |  |
|  |  |  | Pet | 2.29 | (0.5155 - 10.2089) |  |  |  |
|  |  |  | Security | 2.01 | (0.5516 - 7.3294) |  |  |  |
|  | Level of education | No formal education | Primary | 5.03 | (1.5073 - 16.8324) | 16.55 | 4 | 0.002 |
|  |  |  | Secondary | 3.24 | (1.0760 - 9.7699) |  |  |  |
|  |  |  | Tertiary | 7.81 | (2.5085 - 24.3204) |  |  |  |
|  |  |  | Others | 8.5 | (0.4409 - 163.8822) |  |  |  |
| Being bitten by dogs | Occupation | Artisan | Civil Servant | 1.57 | (0.8855 - 2.7869) | 14.40 | 10 | 0.156 |
|  |  |  | Driver | 1.27 | (0.3528 - 4.5395) |  |  |  |
|  |  |  | Farmer | 1.36 | (0.7926 - 2.3460) |  |  |  |
|  |  |  | Fisherman | 3.59 | (0.3168 - 40.581) |  |  |  |
|  |  |  | Laborer | 0.65 | (0.0817 - 5.2051) |  |  |  |
|  |  |  | Security officer | 1.83 | (0.9040 - 3.6966) |  |  |  |
|  |  |  | Student | 2.21 | (0.6814 - 7.1462) |  |  |  |
|  |  |  | Trader/ | 0.68 | (0.3892 - 1.1986) |  |  |  |
|  |  |  | Businessman | 0.98 | (0.6086 - 1.5618) |  |  |  |
|  |  |  | Unemployed | 1.29 | (0.6214 - 2.6915) |  |  |  |

% = Percentage; X<sup>2</sup> = chi-square; DF = Degree of freedom; OR = Odds ratio; 95% CI = 95% confidence interval.

On the contrary, the breed of dogs and the confinement of dogs had a significant effect on the vaccination status of owned dogs. Imported dog breeds were 2.8x (95% CI: 0.7-10.8; p < 0.0001) more likely to be vaccinated than cross-bred or local dogs. Although only 27% (80/293) of dogs were always confined, they are twice more likely (OR: 2.07; 95 CI: 1.2 − 3.7; p = 0.047) to be vaccinated than non-confined dogs. Similarly, the level of education of the respondent’s household had a significant effect on their dog’s vaccination status (p = 0.002). There is a statistically significant difference between the breed of dog and the age at first vaccination with exotic dogs being more likely to be vaccinated at younger ages than local dogs (p = 0.002) (Table S1). In the multivariate analysis fitting all the significant variables affecting the vaccination status (α - 0.25), previous dog-bite victims are 1.7x (95% CI: 0.9530 - 3.1758; p = 0.071) more likely to have vaccinated their dogs against rabies even though previous bite incident remained statistically insignificant. The higher the respondent’s level of education, the more likely they are to vaccinate their dogs (Table S2). Imported breeds of dog were still more likely to be vaccinated than cross-bred or local dogs (Table S2).

## DISCUSSION

Dogs are responsible for 99 of all human rabies (WHO, 2020). Therefore, understanding the domestic dog ecology and its population structure is the first step to an effective elimination program (Cleveland et al., 2006). Baseline epidemiological studies in rabies elimination cannot be overemphasized especially in vaccine procurement for mass vaccination campaigns. More so, they form the evidence-base for the Kwara Rabies Rapid Alert System -KRRAS-; a collaborative state-based project designed to achieve the global aim of eradicating dog-mediated human rabies by 2030 in Kwara state. KRRAS was designed using the five pillars of the global framework; Socio-cultural, Technical, Organization, Political, and Resources (STOP-R) in Kwara state. The total dog population is lower than reported for many other states in Nigeria. This makes Kwara State a good candidate for the first rabies elimination project in the country.

Dog ownership was not affected by religion (p = 0.937). This is in contrast to findings by Mauti et al (2017); Oboegbulem and Nwakonobi (1989) which showed significantly higher dog ownership amongst Christians. Most (85%, n=250/293) dogs were local breeds. This aligns with reports from Kwaghe et al., (2019) and Grace et al., (2018).

Male dogs were more abundant than bitches in Kwara State (sex ratio of dogs 2.2:1). It is believed that male dogs are better guardians and hunters than bitches (Kitala et al., 2001). This is as previously described by Otolorin et al (2014); Aiyedun and Olugasa, (2012). However, this does not agree with the findings of Kwaghe et al., (2019); Hambolu et al (2014) who reported higher bitch to dog ratio. Dogs were mostly kept for security purposes (52%; n=198). This is similar to reports from other parts of Nigeria and Africa (Kwaghe et al., 2019; Garba et al., 2017; Mauti et al., 2017; Otolorin et al., 2014; Kitala et al., 2001). Dog’s age was averaged at 1-3 years. This is similar to reports from all over the country that shows the average dog at > 1 year (Otolorin et al., 2014; Hambolu et al., 2013). Also, Aiyedun and Olugasa (2012) reported 71% of dogs aged > 6 months old.

A dog-human ratio of approximately 1:10 is lower than 1: 7.8 reported in Abia state (Otolorin et al., 2014). Similarly, Kwaghe et al., (2019) reported a higher dog-human ratio of 1: 6.6. Other studies like those of Hambolu et al., (2014); Atuman et. al., (2014); and Garba et al., (2017) reported a dog-human ratio of 1:5.6; 1:4.1 and 1:5.4 respectively (Table 7). In 2012, Magaji (unpublished data) reported a dog-human ratio of 1:28.9 in Kaduna state. Aiyedun and Olugasa (2012) reported a dog-human ratio of 1:139 in Ilorin; which is lower than what this study has recorded in Kwara state. This might be because Aiyedun and Olugasa covered the Ilorin metropolis whilst we studied the whole state. The dog to human ratio reported in this study is within the range reported and modeled for most African countries (Knobel et al., 2005). Within Africa, several studies conducted reported a dog-human ratio of 1:14 in Tanzania (Gsell et al., 2012); 1:21.5 in Chad (Mindekem et al., 2005); 1:15 in Kenya (Kitala et al., 2001) and 1: 16 in Zimbabwe (Brooks R., 1990). Kwara State has a total landmass of 36,825 km^2^, giving a total of 7.5 dogs/ km^2^. This is within the range of 6 and 21 dogs/ km^2^ that was reported by Kitala et al (2001) in Kenya. Much higher dog densities (>1000 dogs/ km^2^) had been estimated for Lagos (Hambolu et al., 2014) and some South American countries (Davlin and Vonville, 2012).

**Table 7.** Comparison of dog ecology across Nigeria and Africa

| Location | Dog-human ratio | Dog density/km<sup>2</sup> | Dog owning households (%) | Dogs/household | Average dog age | Male-Female ratio | Vaccination coverage (%) | References |
| --- | --- | --- | --- | --- | --- | --- | --- | --- |
| **Kwara State (NC)** | 1:10 | 7.5 | 20 | 2.25 | 1-3 years | 2.2:1 | 31 | This study |
| Bauchi (NE) | 1:4.1 | - | - | 2.3 | 1-5 years | 1:1.2 | 26.4 | Atuman et al., 2014 |
| Lagos (SW) | 1:5.6 | - | 95 | 2.8 | > 1 year | 1:1.5 | 64.1 | Hambolu et al., 2014 |
| Abia (SE) | 1:7.8 | - | - | 1.5 | > 1 year | 1.6:1 | 39.9 | Otolorin et al., 2014 |
| Bamako, Mali | 1:121 | 56 | 9 | 0.13 | 3.2 years | 2.8:1 | - | Mauti et al., 2017 |

NC- North-central, NE- North-eastern; SW- South-western; SE-South-eastern Nigeria; % -percentage.

The variations in the dog to human ratio in different areas of Nigeria might be attributable to the different socio-cultural, religious, and economic status of various states of the federation. The vaccination status (31%, n=92/293) is similar to those earlier reported in other parts of the country. This is much lower than the 69.6%; 64.1% and 49.5% reported in Niger state (Garba et al., 2017), Lagos (Hambolu et al., 2014) and Abuja (Mshelbwala et al., 2017) respectively. It is, however, higher than the 26.4% and 21% reported in Bauchi (Atuman et al., 2014) and Nasarawa state (Kwaghe et al., 2019) respectively. This vaccination coverage falls significantly below the 70-80% vaccination rate needed to boost herd immunity (WHO, 2003). With a low vaccination coverage, the general public is at risk of rabies exposure and the need for improved public awareness on rabies, first aid for dog-bite victims, and availability of PEP especially in rural areas cannot be over-emphasized.

Only 26% (n=24/92) of the vaccinated dogs had a vaccination certificate. This is because only licensed veterinarians are allowed to issue a signed vaccination certificate and most vaccinations (52%, n=48/92) were administered at home (possibly- by para veterinary technicians). This has impaired proper monitoring and surveillance of dog health and welfare. The prevalence of dog-bite and its management was evaluated as an important component of rabies epidemiology. The dog-bite incidence was 13% (n=193/1460). This is lower than the 31% and 26.4% reported in Niger and Bauchi states respectively (Garba et al., 2017; Atuman et al., 2014).

There is an urgent need for public education on rabies (mission of KRRAS) as only 15% (n=14/92) of the respondents vaccinated their dogs at the appropriate age of 3-4 months. More so, only 27% (n=61/225) of dog-bite victims collected the post-exposure prophylaxis (PEP) in a health facility. This is especially worrying as most of the respondents (73%) treated dog-bites with antibiotics (13%), non-specific treatment (28%), and traditional (32%) concoctions. Non-specific treatment included wound cleaning and the use of antibiotics. This is highly discouraged as antibiotics have no effect on viruses and it could result in antibiotic resistance. The high level of stray/free-roaming dogs seen in Kwara state is due to the lack of dog population control programs. Hence, the need to re-introduce neutering [or other methods of stray dog population control described under the OIE terrestrial animal health code (OIE, 2019)] during our mass vaccination campaigns. This un-controlled whelping coupled with their use for hunting has introduced and maintained several genera of lyssaviruses in the environment by introducing the sylvatic (wild) rabies cycle into the urban cycle. This could potentially increase the transmission intensity and spread of rabies.

## Conclusion

The results of the study showed that the vaccination coverage needs to be increased, number of dog bites needs to be reduced, number of non-confined dogs need to be reduced, stray dog control strategies need to be intensified, rabies awareness needs to be raised among certain subsets of the population with special emphasis on proper dog-bite treatment regimen and availability of pre-exposure and post-exposure prophylaxis.

This baseline epidemiology and ecology study forms the Phase I and can be used for planning the phase II of KRRAS; the dog Mass Vaccination Campaigns in Kwara state, Nigeria. Information from this study may be used for vaccine procurement, geographical distribution within the state and to serve as a basis for a valid comparison for our post-vaccination surveys. This baseline data is also essential for rabies surveillance in health facilities in Kwara State. We recommend the establishment of a rabies desk office (RDO) and include dog-bite amongst the reportable diseases by the disease surveillance and notification officers (DSNOs) to the District Health Information System (DHIS2). We equally recommend further longitudinal studies to define the health and welfare challenges of dogs in Nigeria.

This is the first comprehensive work describing the dog population structure in Kwara state. The study contributes vital information needed for planning an effective and sustainable rabies control program using the One-Health approach.

## Supporting information

Raw data

## Declarations

### Ethics approval and consent to participate

Ethical clearance was obtained from the various ethical review boards of the Kwara State: Ministry of Health (MoH), Agriculture and Rural Development (MoARD), and Education and Human Capital Development (MoEHCD), Ilorin - Nigeria (reference number: MOH/KS/EU/777/31). Informed consent was sought from the respondents and participants could opt-out at any time.

### Consent for publication

Not applicable

### Availability of data and materials

The survey instrument and datasets are available as supplementary data.

### Authors’ contributions

AIA, AAT, FB, MO, MSO, MSB, GK, and AI were involved in planning the study and data collection. AIA drafted the manuscript. BO and AH did the overall review of the manuscript. All authors read and approved the final study.

## Acknowledgments

We are grateful to the African Field Epidemiology Network (AFENET) for the provision of field data collectors and their server for data upload. We acknowledge the dedication of Jimoh Mohammad, Abdulrahim Ibrahim, and Idris Olarewaju in data collection.

## Supplementary Data

**Table S1:**
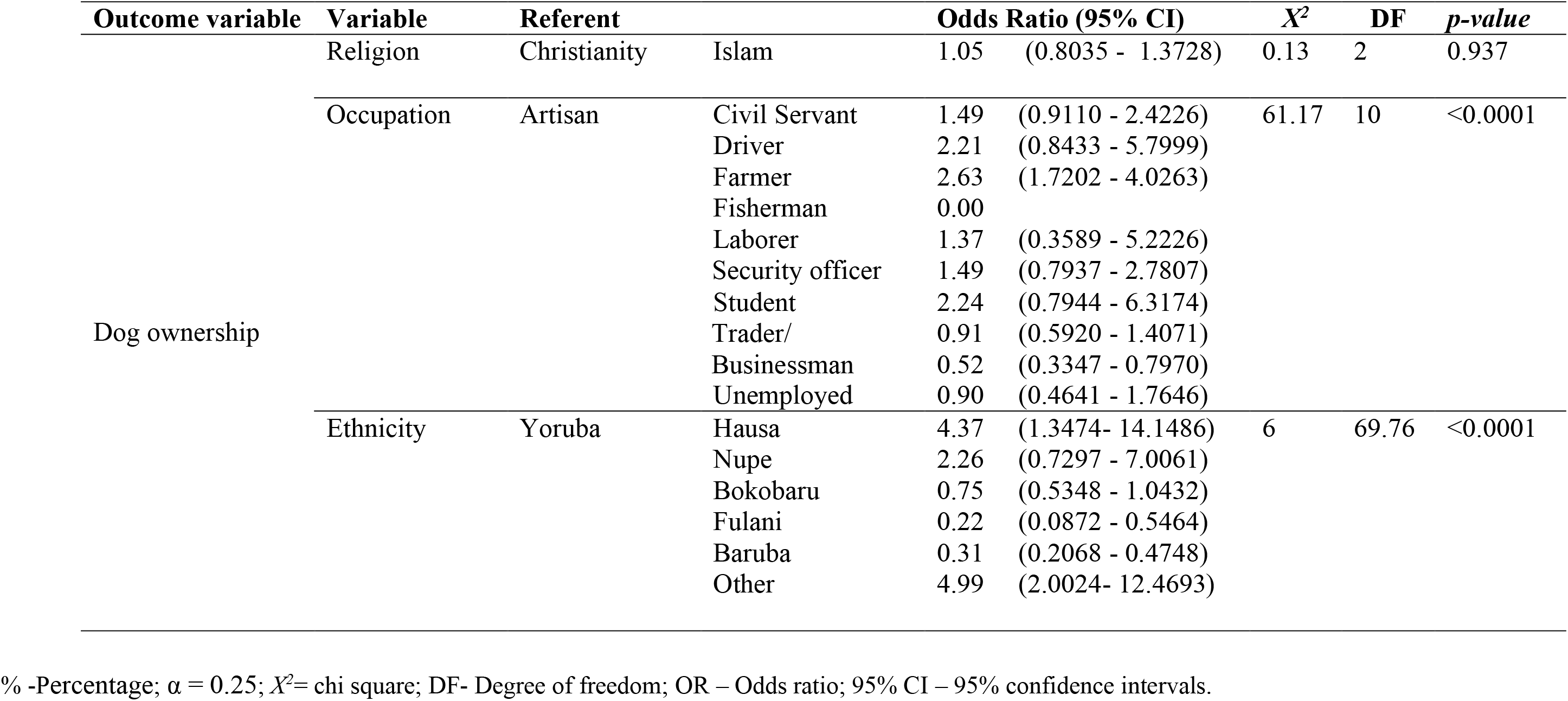
Univariate regression analysis of factors affecting dog-ownership in Kwara state.

**Table S2:** Multivariate analysis of variables affecting dog ownership and the vaccination status of owned dogs in Kwara State.

|  | **Independent Variables** | **Referent** |  | **Adjusted OR (95% CI)** |  | **Adjusted $X^2$** | **DF** | ***p*-value** |
| --- | --- | --- | --- | --- | --- | --- | --- | --- |
| Dog Ownership | Religion | Islam | Christianity | 0.76 | (0.5417 - 1.0644) | 2.55 | 2 | 0.279 |
|  | Ethnicity | Yoruba | Hausa | 3.7673 | (1.1490 - 12.3515) | 69.30 | 6 | 0.000 |
|  |  |  | Nupe | 4.4798 | (1.7716 - 11.3284) |  |  |  |
|  |  |  | Bokobaru | 0.6508 | (0.4423 - 0.9576) |  |  |  |
|  |  |  | Fulani | 0.0431 | (0.0121 - 0.1537) |  |  |  |
|  |  |  | Baruba | 0.2779 | (0.1772 - 0.4358) |  |  |  |
|  |  |  | Other | 2.3748 | (1.1551 - 4.8825) |  |  |  |
| Vaccination Status | Previous dog bite | No | Yes | 1.74 | (0.9530 - 3.1758) | 3.25 | 1 | 0.071 |
|  | Education | No formal education | primary | 5.94 | (1.7298 - 20.4272) | 14.44 | 3 | 0.006 |
|  |  |  | secondary | 3.04 | (0.9736 - 9.4980) |  |  |  |
|  |  |  | Tertiary | 6.99 | (2.1330 - 22.9290) |  |  |  |
|  |  |  | other | 6.15 | (0.2753 - 137.4723) |  |  |  |
|  | Breed | cross | exotic | 2.79 | (0.6482 - 12.0518) | 15.1 | 2 | 0.001 |
|  |  | cross | Local | 0.34 | (0.1375 - 0.8571) |  |  |  |
|  | Gender | Female | Male | 1.47 | (0.8024 - 2.6892) | 1.55 | 1 | 0.213 |

% - Percentage; $\alpha = 0.25$ ; $X^2$ = chi square; DF- Degree of freedom; OR – Odds ratio; 95% CI – 95% confidence intervals.

